# Characterizing Activity and Thermostability of GH5 Cellulase Chimeras from Mesophilic and Thermophilic Parents

**DOI:** 10.1101/382069

**Authors:** Fei Zheng, Josh V. Vermaas, Jie Zheng, Yuan Wang, Tao Tu, Xiaoyu Wang, Xiangming Xie, Bin Yao, Gregg T. Beckham, Huiying Luo

## Abstract

Cellulases from glycoside hydrolase (GH) family 5 are key enzymes in the degradation of diverse polysaccharide substrates and are used in industrial enzyme cocktails to break down biomass. The GH5 family shares a canonical (βα)_8_-barrel structure, where each (βα) module is essential for the enzyme stability and activity. Despite their shared topology, the thermostability of GH5 enzymes can vary significantly, and highly thermostable variants are often sought for industrial applications. Based on a previously characterized thermophilic GH5 cellulase from *Talaromyces emersonii* (*Te*Egl5A, with an optimal temperature of 90°C), we created ten hybrid enzymes with the mesophilic cellulase from *Prosthecium opalus* (*Po*Cel5) to determine which elements are responsible for enhanced thermostability. Five of the expressed hybrid enzymes exhibit enzyme activity. Two of these hybrids exhibited pronounced increases in the temperature optima (10 and 20°C), *T_50_* (15 and 19°C), *T_m_* (16.5 and 22.9°C), and extended half life, *t*_1/2_ (~240- and 650-fold at 55°C) relative to the mesophilic parent enzyme, and demonstrated improved catalytic efficiency on selected substrates. The successful hybridization strategies were validated experimentally in another GH5 cellulase from *Aspergillus nidulans* (*An*Cel5), which demonstrated a similar increase in thermostability. Based on molecular dynamics simulations (MD) of both *Po*Cel5 and *Te*Egl5A parent enzymes as well as their hybrids, we hypothesize that improved hydrophobic packing of the interface between α_2_ and α_3_ is the primary mechanism by which the hybrid enzymes increase their thermostability relative to the mesophilic parent *Po*Cel5.

**IMPORTANCE:** Thermal stability is an essential property of enzymes in many industrial biotechnological applications, as high temperatures improve bioreactor throughput. Many protein engineering approaches, such as rational design and directed evolution, have been employed to improve the thermal properties of mesophilic enzymes. Structure-based recombination has also been used to fuse TIM-barrel fragments and even fragments from unrelated folds, to generate new structures. However, there are not many research on GH5 cellulases. In this study, two GH5 cellulases, which showed TIM-barrel structure, *Po*Cel5 and *Te*Egl5A with different thermal properties were hybridized to study the roles of different (βα) motifs. This work illustrates the role that structure guided recombination can play in helping to identify sequence function relationships within GH5 enzymes by supplementing natural diversity with synthetic diversity.

## INTRODUCTION

Cellulases are a class of industrially important enzymes that have been widely used for biotechnological applications (1, 2). A subset of cellulases isolated from thermophilic microbes are highly thermostable, and display higher cellulolytic activity and half-life at elevated temperatures, which can in turn improve the economic viability of industrial processes by increasing enzymatic hydrolysis rates via operation at higher temperature (3, 4). The GH5 family of endoglucanases is a common component of enzyme cocktails for biomass conversion, and representatives across this family are able to act on diverse oligosaccharide substrates (5). The endoglucanase *Te*Egl5A from *Talaromyces emersonii* exhibits the highest thermostability known so far for this family, with an optimal temperature of 90°C (6). However, this is an outlier among fungal GH5 enzymes, with the most well characterized GH5 enzymes, including those from *Piromyces rhizinflata* (*Pr*EglA) (7), *Thermoascus aurantiacus* (*Ta*Cel5A) (8), *Hypocrea jecorina* (*Trichoderma reesei*) (*Tr*Cel5A) (9), *Ganoderma lucidum* (*Gl*Cel5A) (10), and *Aspergillus niger* (*An*Cel5A) (11) exhibiting an optimal temperature below 70°C. These enzymes are less well suited for some industrial applications, so engineering a thermostable fungal endoglucanase that retains high catalytic activity is desirable.

Various protein engineering approaches, such as rational design and directed evolution, have been employed to improve the thermal properties of mesophilic fungal cellulases (12-14). Among these, SCHEMA, a computational approach to select blocks of sequence with minimal disruption of residue-residue contacts in the resulting functional hybrids (15), offers a useful tool to improve enzyme thermostability. This recombination method has been used to generate novel recombinant enzymes of β-lactamase, β-glucosidases, and GH6 chimeras with significantly higher activity and thermostability (16-18). Structure-based fusion subdomains belonging to different proteins are also an effective method for creating hybrid enzymes with new properties (19).

GH5 enzymes consist of an eightfold repeat of (βα) units that form a barrel, a common (βα)_8_ fold that prior experiments have shown are amenable to improvement through the combinatorial shuffling of polypeptide segments to improve or add functionalities to the protein. For example, the N- and C-terminal four (βα)_4_-barrels of histidine biosynthetic enzymes were assembled to give two highly active hybrid enzymes HisAF and HisFA with the (βα)_8_ fold (20, 21). A high degree of internal structure was exhibited in these two fusion proteins. Sequence-function analysis showed that the recombined protein fragments contributed additively to enzymatic properties in a given chimera (22). In a subsequent experiment, the half barrel of HisF was replaced with (βα)_5_-flavodoxin-like fold from the unrelated but structurally compatible bacterial response regulator CheY, resulting in a stable protein with a (βα)_8_-like fold (23). Later, a catalytically active form of the symmetrical barrel was obtained by fusing two copies of the C-terminal half-barrel HisF-C of HisF (24). Together, these examples suggest that the symmetrical (βα)_8_ barrels like GH5 enzymes are a suitable scaffold for engineering new enzyme properties or functionalities (25).

In this study, we use the plasticity of the classical (βα)8 barrel fold highlighted above to probe the relationship between structure and thermostability in fungal GH5 cellulases by conducting structure-guided protein engineering. The cellulase *Te*Egl5A from *T. emersonii* (6) exhibits high thermostability, retaining almost all of the activity after incubation at 70°C for 1 h, although the structural underpinning for thermal tolerance in this enzyme is unknown. There are many homologous mesophilic proteins. We specifically investigate the cellulase *Po*Cel5 from *Prosthecium opalus* 125034 which retains only <20% activity at 70°C for 10 min. *Po*Cel5 shares the (βα)_8_-barrel structure with *Te*Egl5A along with 51% sequence identity, but shows a much lower temperature optimum, 60°C relative to 90°C, and has been shown to be experimentally tractable to work with (26). By using the fusion approach, the combinations of the first four (βα) module(s) of *Te*Egl5A were introduced into *Po*Cel5, producing ten hybrid enzymes (Table1, Fig. S1). Two of these hybrids exhibit substantial improvements in thermostability relative to the mesophilic parent and catalytic efficiency for specific substrates, which have the potential to lower process costs in industrial bioconversion processes. The functional roles of this N-terminal sequence were also verified in another GH5 cellulase from *Aspergillus niger* (11) and its hybrid. With this work, we determined the structural regions in *Te*Egl5A that contribute to its high thermostability. Comparative molecular dynamics (MD) simulations suggest that improved hydrophobic packing of the interface between α_2_ and α_3_ helices is the primary mechanism behind the improved hybrid thermostability. These simulations also indicate a *Te*Egl5A-specific hydrogen-bond network surrounding R52 that may be an attractive target to further improve GH5 thermostability.

**Table 1:** Schematic structures and molecular masses of the *Po*Cel5-*Te*Egl5A hybrid enzymes^a^

| Protein | Segment(s) | The fragment sequence<br>from *Po*Cel5 | The fragment sequence<br>from *Te*Egl5A | Molecular mass<br>(Da) |
| --- | --- | --- | --- | --- |
| H1 | $\beta_1\alpha_1$ | 47–311 | 1–54 | 35,026 |
| H2 | $\beta_2\alpha_2$ | 1–49/88–311 | 58–95 | 35,445 |
| H3 | $\beta_3\alpha_3$ | 1–89/125–311 | 98–132 | 34,906 |
| H4 | $\beta_4\alpha_4$ | 1–128/160–311 | 137–167 | 34,921 |
| H5 | $\beta_1\alpha_1+\beta_2\alpha_2$ | 88–311 | 1–95 | 35,605 |
| H6 | $\beta_2\alpha_2+\beta_3\alpha_3$ | 1–49/125–311 | 58–132 | 34,921 |
| H7 | $\beta_3\alpha_3+\beta_4\alpha_4$ | 1–89/160–311 | 98–167 | 34,801 |
| H8 | $\beta_1\alpha_1+\beta_2\alpha_2+\beta_3\alpha_3$ | 125–311 | 1–132 | 35,485 |
| H9 | $\beta_1\alpha_1+\beta_2\alpha_2+\beta_3\alpha_3+\beta_4\alpha_4$ | 160–311 | 1–167 | 35,381 |
| H10 | $\beta_5\alpha_5+\beta_6\alpha_6+\beta_7\alpha_7+\beta_8\alpha_8$ | 1–159 | 168–317 | 34,843 |

<sup>a</sup> Residue numbering relative to the parent sequences

## RESULTS

### Cloning and sequence analysis of *Po*Cel5

A gene fragment, 341 bp in length, was amplified from the genomic DNA of *P. opalus* CBS 125034, using GH5-specific primers. The 5′ and 3′ flanking regions were obtained by TAIL-PCR and assembled with the known sequence to give full-length *PoCel5* (1062 bp). Sequence analysis indicated that the ORF of *Po*Cel5 is interrupted by one intron (69 bp). The cDNA of *PoCel5* contained 993 bp that encoded a GH5 endoglucanse of 330 amino acids with an estimated molecular mass of 35 kDa and a predicted *p*I value of 4.94. Based on high sequence homology with other GH5 cellulases of known structure, *Po*Cel5 likely contains only one catalytic domain with a (βα)_8_-barrel fold. Similarly, N-terminal 19 amino acids were predicted to be a signal peptide, and three N-linked glycosylation sites (Asn23, Asn64, and Asn76) are possible based on protein sequence and structure (Fig. 1).

**FIGURE 1.**
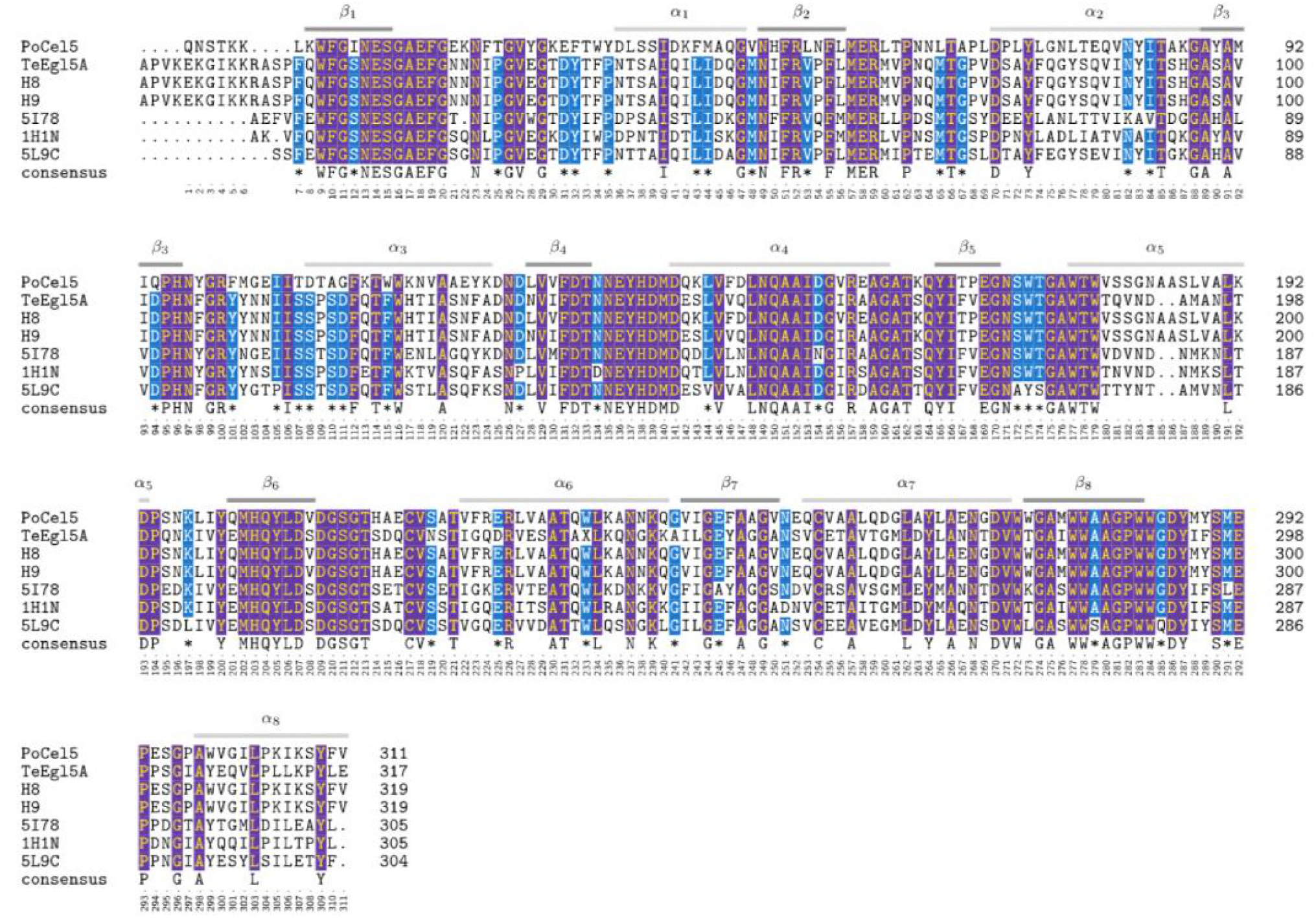
Sequence alignment of *Po*Cel5, *Te*Egl5A, and their hybrid enzymes H8 and H9 and three crystallized GH5 cellulases, 5I78 from *Aspergillus niger* (11), 1H1N from *Thermoascus aurantiacus* (59), and 5L9C from *Penicillium verruculosum*. Residue numbering at the bottom is numbered according to the sequence from *Po*Cel5, which is used consistently throughout the text when referring to residue numbers, including for *Te*Egl5A and the hybrids to simplify interaction comparison between enzymes. Module elements are labelled above the sequence. Residues highlighted in purple are identical across all sequences, with residues highlighted in blue being conserved in all but 1 sequence.

### Design and production of a chimeric enzyme library

The thermophilic *Te*Egl5A enzyme (temperature optimum at 90°C), exhibits high catalytic efficiency and broad substrate specificity (6), and shares a common (βα)_8_-barrel fold with *Po*Cel5. This (βα)_8_-TIM-barrel structure is common in 28 GH families, which include xylanases, cellulases, mannanases, amylases, and triosephosphate isomerases (CAZy; http://www.cazy.org) (27), and may have evolved through gene duplication event as in a previously studied imidazole glycerol phosphate synthase (28). Catalytic amino acids typical of GH5 cellulases are also shared between both *Te*Egl5A and *Po*Cel5.

Based on this common structure and functionality, we swapped sequence fragments between *Te*Egl5A and *Po*Cel5 to create a chimeric cellulase library. These sequence swaps occur at the boundary of individual (βα) modules. Within each module, there is a β-strand and α-helix linked together by a βα-loop. For enzymes with a (βα)_8_ topology, the eight β-strands assemble into a central β-sheet, i.e. the barrel, which is surrounded by eight α-helices (10). Guided by the enzyme structures, the N-terminal and C-terminal modules of endoglucanases *Te*Egl5A, *Po*Cel5, and *An*Cel5A were used to design twelve fusion proteins, combinatorically isolating the specific fragments that result in improved thermostability (Fig. S1 and Table 1).

### Expression and purification of *Po*Cel5 and hybrid enzymes

Recombinant *Po*Cel5 was successfully produced in *P. pastoris* GS115 component cells after a 48-h methanol induction. The enzyme was purified to electrophoretic homogeneity (Fig. S2a) but with an apparent molecular mass higher than the predicted value (35 kDa), with glycosylation likely accounting for the higher molecular weight. After treatment with Endo H to remove glycosylation, recombinant *Po*Cel5 migrated as a protein band corresponding to its expected molecular mass.

Five (H4, H5, H6, H8, and H9) of the ten *Po*Cel5-*Te*Egl5A hybrid enzymes exhibited cellulase activity measured by DNS. In comparison to the wild-type, these active hybrids showed various protein migration patterns in the gel before and after Endo H treatment to remove enzyme glycosylation (Fig. S2a). The different numbers of structure-based *N*-glycosylation sites, four (Asn2, Asn23, Asn63, and Asn76) for H4, two (Asn21 and Asn44) for H5, two (Asn2 and Asn23) for H6, one (Asn36) for enzymes H8, and H9, may account for this variation. After the Endo H treatment, all hybrid enzymes exhibited a molecular mass equal to their estimated values (Table 1).

### Effect of pH

The effect of pH on the activity and stability of wild-type and hybrid enzymes was determined using CMC-Na as the substrate. The wild-type and hybrid H4 were optimally active at pH 5.0, while the others had a pH optimum at 4.0 (Fig. 2a). The enzymes exhibited pH-dependent stability (Fig. 2b). The wild-type and hybrids H4–H6 retained long-term activity over a pH range of 4.0 to 7.0 (>80% activity), were largely ineffective and possibly unfolded at pH 8.0–9.0, but retained >50% activity at pH 10.0–12.0. In contrast, hybrids H8 and H9 had a broader pH stability range, retaining >70% activity at pH 3.0 to 10.0.

**FIGURE 2.**
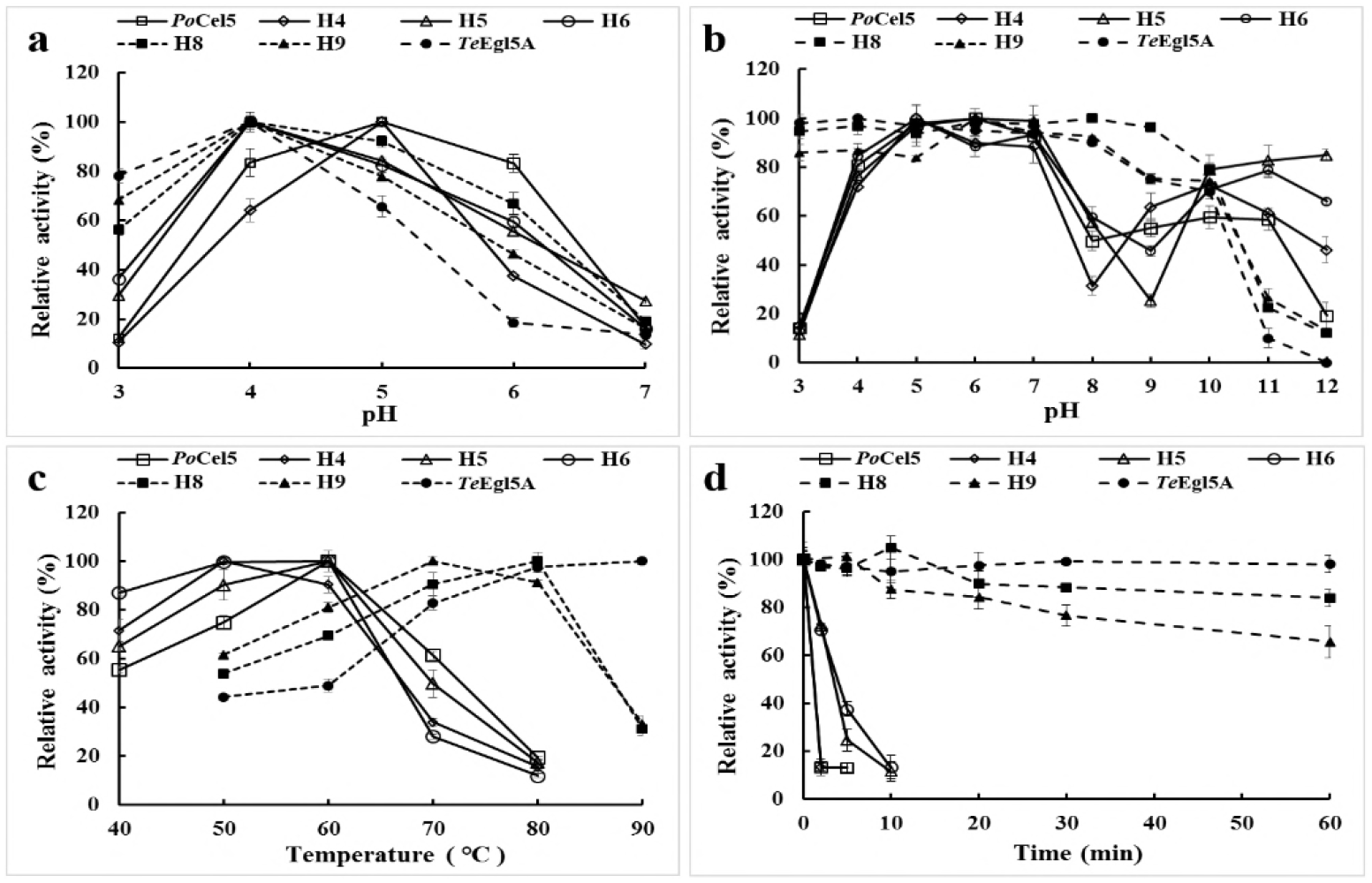
Enzyme properties of the purified recombinant *Po*Cel5, TeCel5 and their hybrid enzymes. The relative activities corresponding to 100% are 351 U/mg for *Po*Cel5, 620 U/mg for *Te*Egl5A, 270 U/mg for H4, 175 U/mg for H5, 101 U/mg for H6, 721 U/mg for H8, and 917 U/mg for H9, respectively, with CMC-Na as the substrate. (a) pH-activity profiles tested at the optimal temperature of each enzyme (60°C for *Po*Cel5, 50°C for H4, H5 and H6, 80°C for H8, 70°C for H9, and 90°C for *Te*Egl5A). (b) pH-stability profiles. After incubation of the enzymes at 37°C for 1 h in buffers ranging from pH 3.0 to 12.0, the residual activities were determined in 100 mM citric acid-Na_2_HPO_4_ buffer at optimal pH and optimal temperature of each enzyme. (c) Temperature-activity profiles tested at the optimal pH of each enzyme (pH 5.0 for *Po*Cel5 and H4, and pH 4.0 for *Te*Egl5A, H5, H6, H8, and H9). (d) Temperature-stability profiles. Each enzyme was pre-incubated at 70°C and optimal pH in 100 mM citric acid-Na_2_HPO_4_ buffer for different periods of time, and subjected to residual activity assay under optimal conditions of each enzyme.

### Thermal property analysis

To determine the optimal temperature of *Po*Cel5 and hybrids, their enzyme activities at different temperatures (40–90°C) were determined after a 10 min reaction with 1% CMC-Na in 100 mM citric acid-Na_2_HPO_4_ at optimal pH. The optimal temperature of wild-type *Po*Cel5 was determined to be 60°C (Fig. 2c). When replacing the (βα) module(s) of *Po*Cel5 with those from *Te*Egl5A, the temperature optima of all hybrids were lower than the 90°C for *Te*Egl5A. H4 exhibited maximum activity at 50°C, H5 and H6 had a temperature optimum of 60°C similar to the wild-type, and H8 and H9 showed optimal activities at 80°C and 70°C, respectively. Significant differences were also seen in their thermostability (Fig. 2d). When incubated at 70°C, *Po*Cel5 as well as hybrids H4–H6 lost activity rapidly, retaining <20% activity within 10 min. By contrast, H8 and H9 retained >60% activity for 1 h, indicating that in this case the replacement of the N-terminal (βα) module(s) from the mesophilic *Po*Cel5 with the thermophilic *Te*Egl5A improved hybrid thermostability.

The thermal stability parameters of the wild-type *Po*Cel5 and *Po*Cel5-*Te*Egl5A hybrids were also compared (Table 2). The *T*_50_, *T*_m_, and *t*_1/2_ values at 55°C of *Po*Cel5 were 57°C, 53.6°C, and 0.4 h, respectively. In comparison to the wild-type, hybrids H6, H8, and H9 show higher *T*_50_ (7–19°C increase) and *T*_m_ (7.6–22.9°C increase) values and longer *t*_1/2_ (16–650-fold) at 55°C, with H8 and H9 being more thermostable.

**Table 2:** Thermodynamic properties of *Po*Cel5 and its hybrid enzymes

| Enzyme | $T_{50}$ (°C) | $T_m$ (°C) | $t_{1/2}(55^\circ\text{C})$ (h) |
| --- | --- | --- | --- |
| *Po*Cel5 | $57 \pm 1$ | $53.6 \pm 1.2$ | $0.40 \pm 0.05$ |
| H4 | $58 \pm 2$ | $51.2 \pm 1.0$ | $0.06 \pm 0.01$ |
| H5 | $56 \pm 1$ | $49.3 \pm 1.1$ | $0.50 \pm 0.04$ |
| H6 | $64 \pm 1$ | $61.2 \pm 1.0$ | $6.5 \pm 2.5$ |
| H8 | $76 \pm 2$ | $76.5 \pm 1.3$ | $260 \pm 3.5$ |
| H9 | $72 \pm 2$ | $70.1 \pm 1.2$ | $96 \pm 1.5$ |

### Specific activity and kinetics

GH5 cellulases can often act on multiple substrates, including glucan substrates as well as birchwood xylan, Avicel, and laminarin (6, 29, 30). Table 3 shows the activity of the enzyme library on four substrates. Enzymatic digestion was fastest for barley β-glucan, lichenan, and CMC-Na, with comparatively slow activity when given locust bean gum as a substrate. The enzymes demonstrated no activity with birchwood xylan, Avicel, and laminarin, suggesting specificity typical of cellulolytic endoglucanases. The specific activities of *Po*Cel5 against barley β-glucan, lichenan, and CMC-Na were higher than that of hybrids H4–H6 with the replacements of β_4_α_4_ (H4), β_1_α_1_ + β_2_α_2_ (H5), and β_2_α_2_ + β_3_α_3_ (H6), but much lower than that of hybrids H8 and H9 and *Te*Egl5A. These results indicate that introduction of the N-terminal three (hybrid H8) and four (hybrid H9) blocks of (βα) modules of *Te*Egl5A improved the catalytic performance of *Po*Cel5. H9 exhibited the highest specific activities among all enzymes against barley β-glucan, lichenan, and CMC-Na, which were 20%, 116%, and 160% higher than that of the parental *Po*Cel5 enzyme, respectively. The improved catalysis is in some cases non-additive, as both H8 and H9 demonstrated improved performance on CMC-Na relative to both parent enzymes.

**Table 3:** Specific activities of *Po*Cel5, *Te*Egl5A and their hybrid enzymes

| Enzymes | Barly β-glucan<br>(U/mg) | Lichenan<br>(U/mg) | CMC-Na<br>(U/mg) | Locust bean gum<br>(U/mg) |
| --- | --- | --- | --- | --- |
| *PoCel5* | 1535 ± 13 | 1436 ± 81 | 351 ± 6 | 82 ± 6 |
| *TeEgl5A* | 2688 ± 23 | 3291 ± 34 | 596 ± 26 | 96 ± 13 |
| H4 | 1130 ± 81 | 1424 ± 72 | 270 ± 11 | 71 ± 1 |
| H5 | 549 ± 16 | 633 ± 33 | 175 ± 2 | 30 ± 2 |
| H6 | 103 ± 2 | 159 ± 19 | 101 ± 1 | 1.4 ± 0.1 |
| H8 | 1606 ± 51 | 2548 ± 46 | 722 ± 30 | 55 ± 3 |
| H9 | 1844 ± 87 | 3107 ± 122 | 917 ± 50 | 74 ± 6 |

To better understand the effect of different motifs from *Te*Egl5A on the catalytic performance of *Po*Cel5, kinetic studies of the wild-type *Po*Cel5 and *Po*Cel5-*Te*Egl5A hybrids were performed using CMC-Na as the substrate at the optimal reaction conditions of each enzyme (Table 4). The *K*_m_ and *V*_max_ values of *Po*Cel5 were 4.9 ± 0.3 mg/mL and 647 ± 46.7 mol/min/mg, respectively, and its catalytic efficiency was *k*_cat_/*K*_m_ 76.2 ± 1.8 mL/s/mg, which is lower than that of the EG (118 mL/s/mg) from *Penicillium purpurogenum* (31). The kinetic values of the hybrids had the same trends as the specific activities. In comparison to the *Po*Cel5 parent, hybrids H4–H6 showed decreased substrate affinity (higher *K*_m_ values) and reduced reaction velocity and turnover rates (lower *V*_max_ and *k*_cat_ values), while hybrids H8 and H9 showed greater substrate affinity, reaction velocity, and turnover rates. The catalytic efficiencies of the hybrids H4–H6 decreased to 17.4–51.4% to that of *Po*Cel5, and those of H8 and H9 increased up to 276%. The H8 and H9 enzymes specifically are in some sense even more efficient than the thermophilic *Te*Egl5A parent, as they combine the high affinity of *Po*Cel5 with the higher turnover rate of *Te*Egl5A for a 2-3 fold improvement in catalytic efficiency (*k*_cat_/*K*_m_) relative to either parental enzyme. These findings indicate that the replacement of different (βα) modules from *Te*Egl5A could either enhance or prove deleterious to the function of *Po*Cel5, depending on the specific replacements made. The primary differences in the five *Po*Cel5-*Te*Egl5A hybrids in catalytic performance (substrate binding, enzyme catalysis, and product dissociation) may be attributed to their structural differences. The hybrids H4–H6 had only one or two (βα) module(s) from *Te*Egl5A, while the hybrids H8 and H9 contained three or four (βα) exogenous modules (Table 1). Hybrids H5 and H8 had one module difference (β_1_α_1_+β_2_α_2_ vs. β_1_α_1_+ β_2_α_2_+β_3_α_3_), but varied in temperature optima (60°C vs. 80°C), thermal stability, and catalytic performance. Based on these results alone, we conjecture that the β_3_α_3_ module may play a key role in protein structure and catalysis. However, hybrid H3 (containing the β_3_α_3_ of *Te*Egl5A alone) did not show improvements in the properties examined in this study. Moreover, the hybrids H4 containing partial segments of the replaced modules of H8 and H9 had no improvement in stability and catalysis.

**Table 4:**
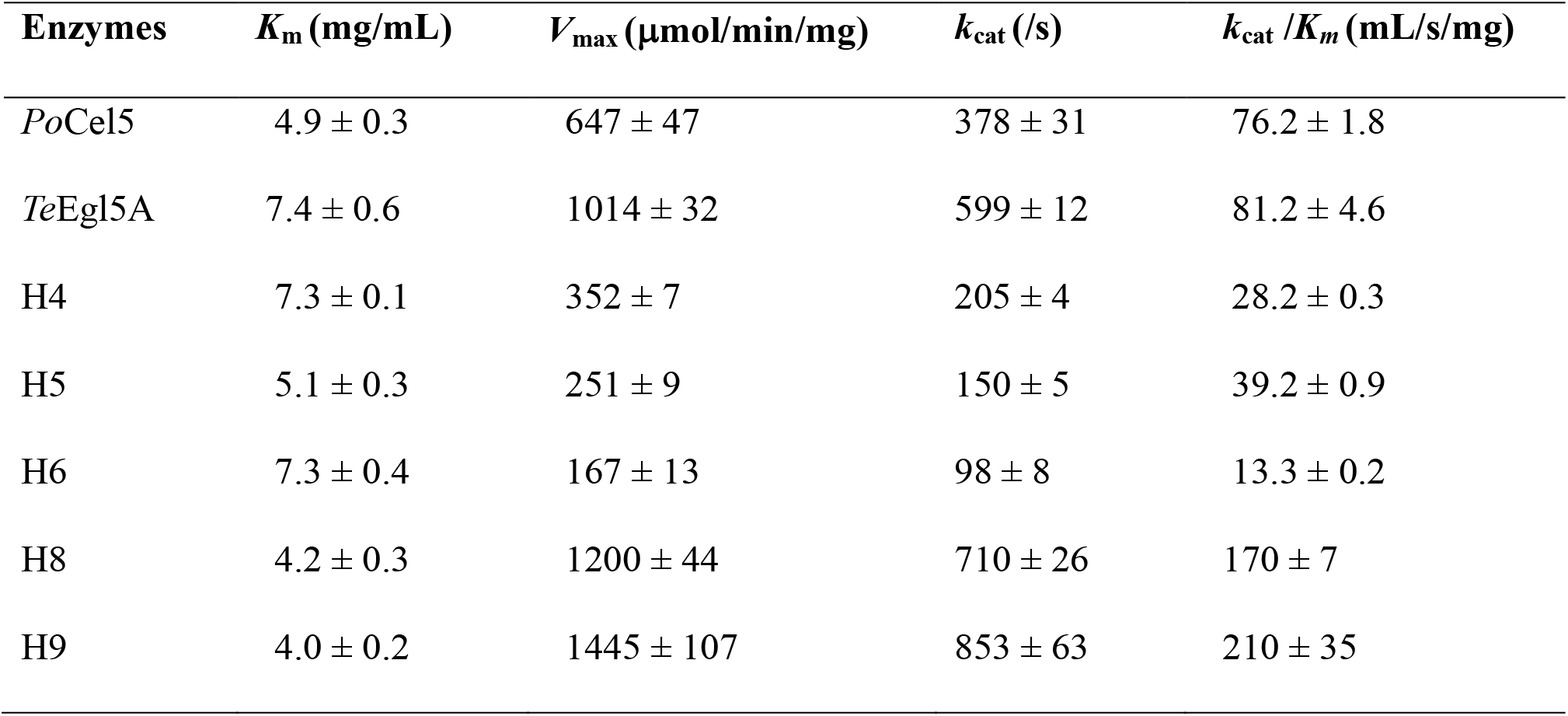
Kinetic parameters of *Po*Cel5, *Te*Egl5A and their hybrid enzymes with CMC-Na as the substrate

### Functional verification by *An*Cel5 and its hybrids

To verify the common effect of the (βα)_1-3_ and (βα)_1-4_ modules of *Te*Egl5A, the corresponding part of another GH5 cellulase from *A. nidulans* was also replaced (11). Two hybrid enzymes (*An*Cel5-H1 and *An*Cel5-H2) were constructed and produced in *P. pastoris* GS115, but only *An*Cel5-H2 showed cellulase activity. After purification to homogeneity (Fig. S2b), the wild-type *An*Cel5 and hybrid *An*Cel5-H2 were biochemically characterized. Both *An*Cel5 and *An*Cel5-H2 were optimally active at pH 4.0 (Fig. S3a). However, *An*Cel5-H2 showed adaptability and stability over higher temperatures. It had an optimal temperature of 80°C, 10°C higher than that of the wild-type *An*Cel5 (Fig. S3b). *An*Cel5 was stable at temperature ≤60°C, while the hybrid *An*Cel5-H2 retained stability at 70°C (Fig. S3c). Moreover, *An*Cel5 lost activity rapidly at 70°C and 80°C (retaining <10% within 20 min); under the same conditions, *An*Cel5-H2 retained >30% activity after a 1 h-incubation (Fig. S3d). The *T*_50_, *T*_m_, and *t*_1/2_ (70°C) values of *An*Cel5 and *An*Cel5-H2 were 70 ± 2 and 78 ± 1°C, 83.5 ± 1.0 and 87.0 ± 1.0°C, and 0.6 ± 0.1 and 9.0 ± 0.6 h, respectively. When using CMC-Na as the substrate, *An*Cel5 and *An*Cel5-H2 showed a specific activity of 1818 ± 26 and 2800 ± 34 U/mg respectively, similar *V*_max_ (2397 ± 57 vs. 2486 ± 36 μmol/min/mg) and *k*_cat_ (1402 ± 35 vs. 1462 ± 47 /s) values, but different *K*_m_ values (4.2 ± 0.3 vs. 3.4 ± 0.2 mg/mL). When combined with the previous results, selective recombination of different related enzymes can indeed be an effective strategy for improving the thermal stability and catalytic efficiency of GH5 fungal cellulases.

### Mechanism of improved thermostability

The evidence presented indicates that there are structural elements unique to the N-terminal region of the thermophilic *Te*Egl5A that enhance the performance of the enzyme at high temperature. Based on the determined melting temperatures (Table 2), a region of particular interest would be the interface between β_2_α_2_ and β_3_α_3_, which when replaced by the *Te*Egl5A equivalent as in H6, raises the melting temperature relative to unaltered *Po*Cel5 by nearly 10°C. Additional 10–15°C increases in the melting temperature when β_1_α_1_ from *Te*Egl5A are also included, as in H8 and H9, but not when independently replaced, as in H1 and H5, suggest that the β_1_α_1_ module also is involved in the denaturation process of these hybrid enzymes.

To explore which specific interactions may participate in the increased melting points of specific chimeras, conventional equilibrium MD simulations were performed for models of H8, H9, as well as both parent enzymes. By varying the simulation temperature from 25°C (298K) to near the melting point 70°C (343K) and beyond to 125°C (398K), we track how these interactions change with temperature over the course of the five replicates for each combination of temperature and protein. By correlating specific interactions with temperature change across the conformational ensemble created by simulation, we resolve the improved hydrophobic packing at the interface between modules 2 and 3. Since single simulations are only 200 ns long, only minimal thermal denaturation is observed. Together, the results from the MD simulations provide mechanistic insight as to the denaturation process, and how it is arrested in part in the H8 and H9 hybrids.

### Barrel stabilization by hydrogen bond networks

A commonly invoked mechanism to improve protein thermostability are rigidifying mutations (32), which would result in significantly lower fluctuations for specific residues. Using MD simulations, we can directly test for reduced fluctuations at specific residue positions by computing the root mean squared fluctuation (RMSF), which measures the mean fluctuation away from the average position of that residue (Fig. 3). We find that there is a consistent reduction in fluctuation at elevated temperature only for the original thermophilic *Te*Egl5A, particularly in structured regions within the β-sheets. The hybrid enzymes, by contrast, have fluctuations in line with what was observed in *Po*Cel5, even in regions where the sequence was identical to *Te*Egl5A (β_1_α_1_-β_3_α_3_ for H8, β_1_α_1_-β_4_α_4_ for H9). Thus, the increased thermostability of H8 and H9 are not strictly due to rigidifying mutations.

**FIGURE 3.**
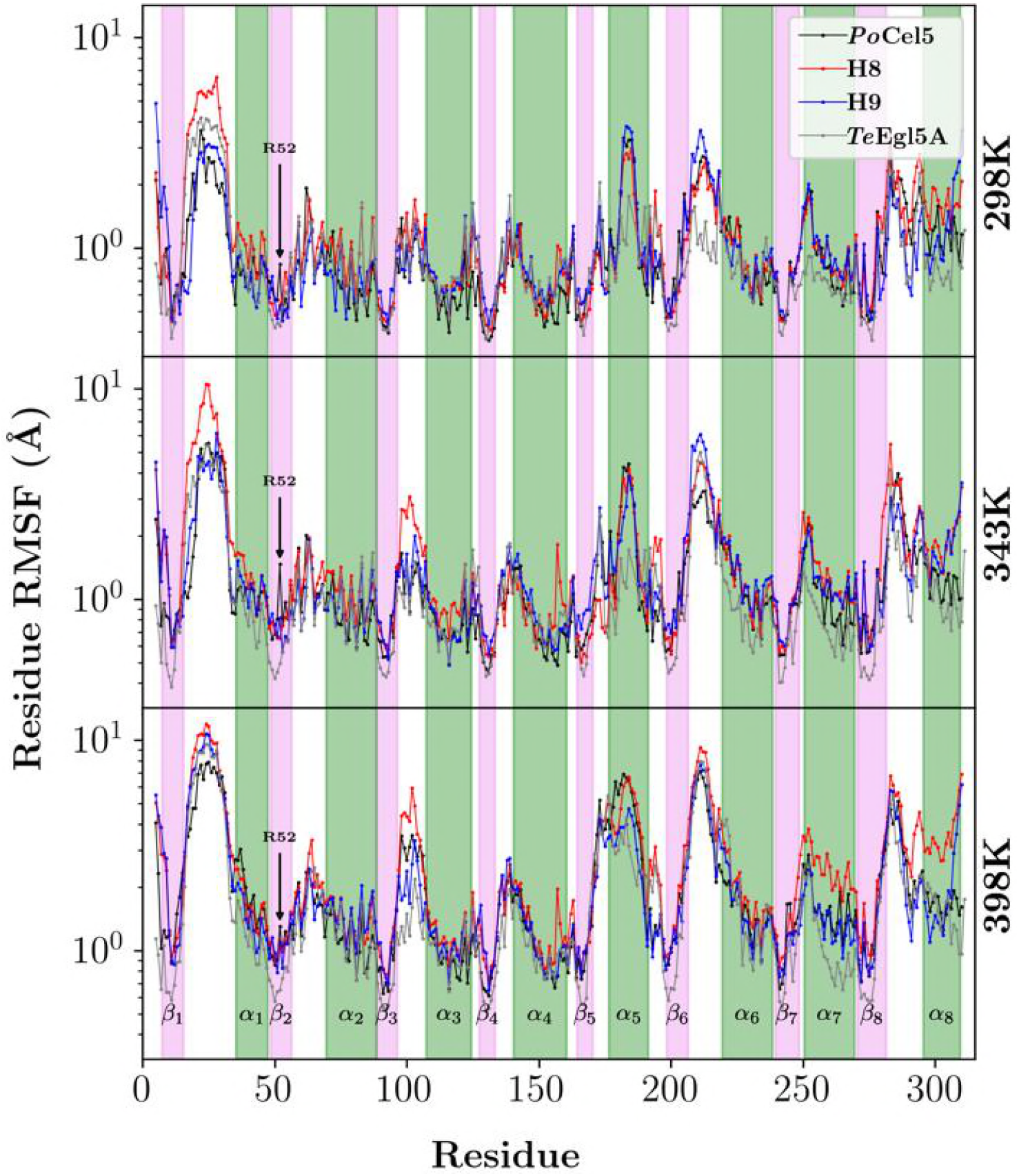
Comparative per residue root mean square fluctuation (RMSF) across all trajectories. The RMSF for each simulated GH5 is presented on three subgraphs (*Po*Cel5 black, H8 red, H9 blue, and *Te*Egl5A gray), one for each temperature as indicated. To highlight the disparity of RMSF between structural elements and the intervening loops, the eight beta strand regions are highlighted in pink and the eight alpha-helical regions are highlighted in green, as well as labelled in the 398K subpanel. The elevated RMSF for *Po*Cel5 at residue 52 is indicated by a black arrow.

Instead, the reduced RMSF for the β-sheets within *Te*Egl5A imply that there are specific interactions formed within the barrel of *Te*Egl5A that are not present in the *Po*Cel5 or its hybrids. From the sequences alone, it is not clear which residues are in close proximity and might interact across the barrel. *Te*Egl5A sequence introduces a number of charged residues within the barrel relative to *Po*Cel5, some of which were determined to be protonated at physiological pH by pKa estimation tools (33) based on their inaccessibility to water and possible hydrogen bonds that could be made with neighboring residues. The network of interactions that is formed in *Te*Egl5A (Fig. 4B) directly connects together more structural elements relative to the interactions seen in *Po*Cel5 (Fig. 4A), reducing the fluctuation increases within the central barrel as the temperature rises (Fig. 3). In the hybrid enzymes, only some of these interactions are retained. There is effectively a Q94D mutation in both H8 and H9 due to the sequence they inherit from *Te*Egl5A. This change relative to *Po*Cel5 is sufficient to provide R52 a strong binding partner and significantly lower the fluctuations of the R52 residue in the hybrids, as indicated by the black arrows in Fig. 3.

**FIGURE 4.**
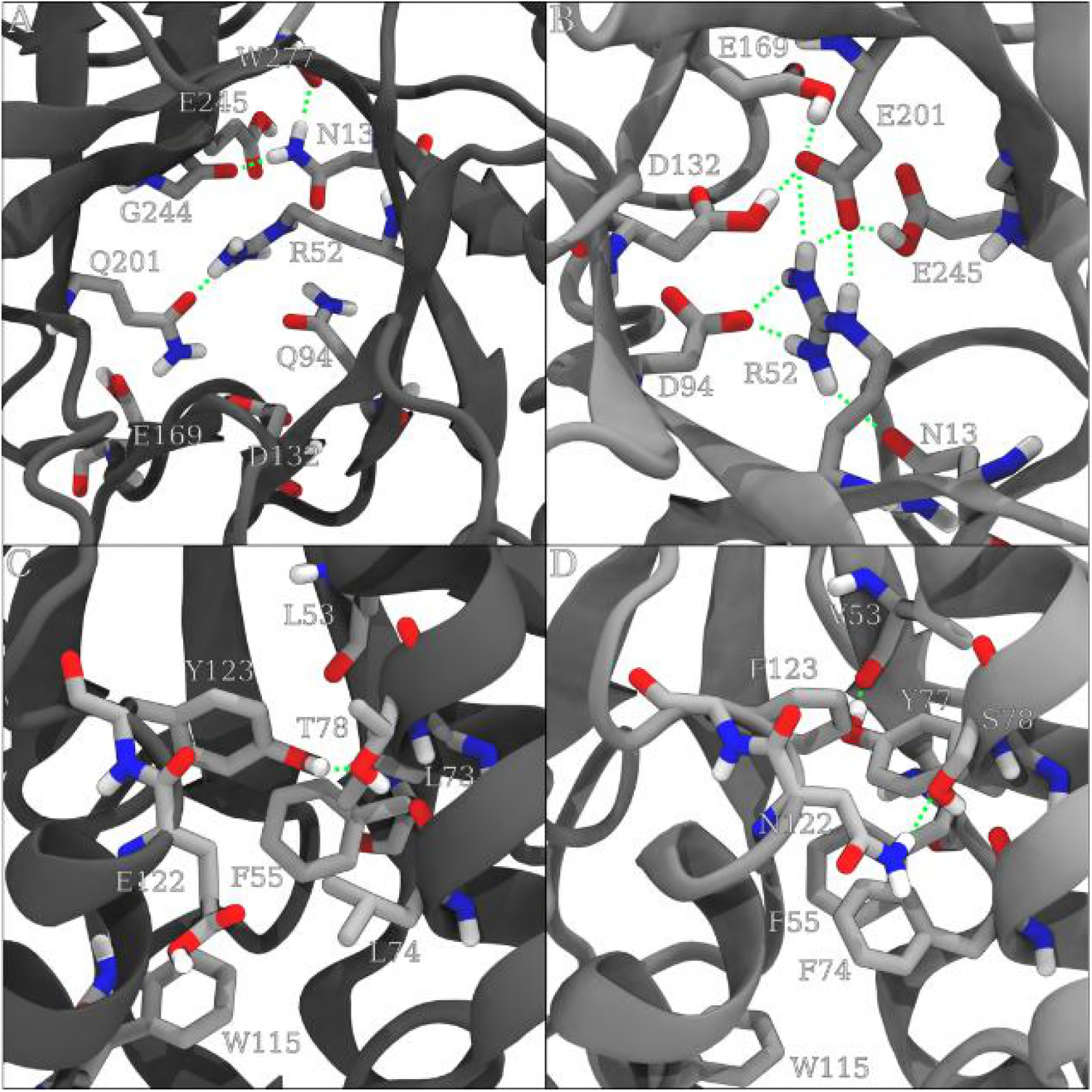
Simulation snapshots. In the barrel assembly of *Po*Cel5 (A) and *Te*Egl5A (B), we highlight specific interactions around the R52 residue present for each enzyme. For visual clarity, only heavy atoms and polar hydrogens are shown for the selected residues that form an extended hydrogen bond network with R52. Along the interface between β_2_α_2_β_3_α_3_, sequence differences between *Po*Cel5 (C) and *Te*Egl5A (D) lead to different hydrogen bonds and hydrophobic packing arrangements. The hydrogen bond interactions are shown as green dashed lines.

Other interactions found in *Te*Egl5A that further stabilize the central barrel are missing from the hybrids. One example occurs at position 201, where H8 and H9 retain Q201 instead of E201 found in *Te*Egl5A. As a result, neighboring charged residues do not form extended hydrogen bonding networks as they do in *Te*Egl5A (Fig. 4B), and instead show a lack of interaction similar to *Po*Cel5 (Fig. 4A). At low temperatures, the weaker hydrogen bond networks formed with N13 and sporadic interactions between Q94 and Q201 are sufficient to stabilize the barrel. However, without the central barrel stabilization brought about by the additional hydrogen bonds seen in *Te*Egl5A, the barrel exhibits higher fluctuations with increasing temperature. The extensive *Te*Egl5A interaction network increases thermostability of the barrel complex, and may be an additional avenue by which thermostability could be further improved, similar to efforts in other fungal cellulases to add hydrogen bonds to improve thermostability (34, 35).

### Specific Interactions in β_2_α_2_ and β_3_α_3_

To narrow down what subdomain components are interacting, we first evaluated hydrogen bonds within the first 4 module sets (Table 5). In this analysis, we see that *Po*Cel5 actually creates hydrogen bonds more frequently within the first 4 modules than do the hybrids or the thermophilic enzyme. This difference shrinks when the same analysis is conducted on the higher temperature simulations (Tables S2 and S3), indicating that these interactions are not as stable overall. In part, this may be due to the unexpected hydrogen bond formed between Y77 and V53 in the *Te*Egl5A and the hybrids H8 and H9 (Fig. 4D) but is not present in *Po*Cel5 (Fig. 4C). This interaction is the only direct hydrogen bond formed that goes between a helical segment and a beta strand within the first four modules of the enzyme (Table 5). When combined with the adjacent rigidifying interactions in the barrel core surrounding R52, this provides part of a mechanism of how sequence replacement improves thermostability.

**Table 5:** Hydrogen bond propensities at 298K across all simulations for residues within the β_1-4_ or α1-4 modules, excluding hydrogen bonds involved in helical interactions. A hydrogen bond is counted only if the heavy atoms are within 3.2 Angstrom and the hydrogen is no more than 30 degrees removed from the direct line between the heavy atoms. To help quickly identify the structural element each residue is found in, the residue identifiers are color coded. Module 1 uses black, module 2 uses blue, module 3 uses green, and module 4 uses red. The lightness of the color indicates whether it is in a β element (dark) or α (lighter).

| *PoCel5* |  | H8 |  | H9 |  | *TeEgl5A* |  |
| --- | --- | --- | --- | --- | --- | --- | --- |
| Donor/Acceptor | H-bond | Donor/Acceptor | H-bond | Donor/Acceptor | H-bond | Donor/Acceptor | H-bond |
| M88/P47 | 0.93 | R48/D90 | 1.17 | R48/D90 | 1.39 | V88/P47 | 0.95 |
| L49/M88 | 0.91 | D128/D90 | 0.98 | V88/P47 | 0.89 | R48/D90 | 0.92 |
| K113/E154 | 0.86 | V88/P47 | 0.94 | I89/I126 | 0.82 | V49/V88 | 0.87 |
| V126/A87 | 0.85 | V49/V88 | 0.88 | I126/A87 | 0.80 | I89/I126 | 0.87 |
| Q90/L49 | 0.85 | I89/V126 | 0.86 | S86/N45 | 0.80 | P6/N45 | 0.87 |
| K109/D150 | 0.84 | P6/N45 | 0.84 | A87/N124 | 0.76 | E10/R48 | 0.81 |
| Y86/N45 | 0.79 | V126/A87 | 0.83 | E10/R48 | 0.75 | P47/S86 | 0.79 |
| E10/R48 | 0.78 | S86/N45 | 0.75 | D128/D90 | 0.74 | I126/A87 | 0.77 |
| K139/D143 | 0.76 | E10/R48 | 0.73 | V49/V88 | 0.72 | S86/N45 | 0.77 |
| A87/L124 | 0.76 | A87/L124 | 0.72 | P51/D90 | 0.66 | A87/N124 | 0.70 |
| P6/N45 | 0.74 | P47/S86 | 0.71 | Y73/V49 | 0.65 | P51/D90 | 0.68 |
| K109/E154 | 0.73 | P51/D90 | 0.67 | P47/S86 | 0.65 | D107/S104 | 0.65 |
| R48/I8 | 0.71 | Y73/V49 | 0.67 | R48/S8 | 0.63 | R48/N9 | 0.64 |
| R48/N9 | 0.69 | K139/D143 | 0.64 | P6/N45 | 0.63 | D128/I89 | 0.63 |
| P51/Q90 | 0.68 | N45/Q4 | 0.60 | N45/P6 | 0.56 | N45/P6 | 0.53 |
| Y119/T74 | 0.65 | R48/N9 | 0.60 | D128/I89 | 0.47 | Y73/V49 | 0.52 |
| S34/D32 | 0.59 | N45/P6 | 0.60 | Y79/Q37 | 0.41 | R48/S8 | 0.37 |
| Q90/N50 | 0.58 | R48/S8 | 0.58 | D107/S104 | 0.40 | A85/I80 | 0.27 |
| N45/P6 | 0.57 | R48/D128 | 0.52 | N45/Q4 | 0.39 | S8/N9 | 0.27 |
| G107/D104 | 0.56 | D128/I89 | 0.45 | S8/N9 | 0.32 | N45/Q4 | 0.26 |
| N50/E10 | 0.54 | S104/D107 | 0.39 | A85/I80 | 0.30 | S117/T114 | 0.26 |
| K83/D37 | 0.53 | A85/I80 | 0.32 | S104/D107 | 0.29 | P119/A116 | 0.21 |
| P47/Y86 | 0.52 | D107/S104 | 0.31 | R48/N9 | 0.24 | Y79/Q37 | 0.20 |
| N45/K4 | 0.51 | K139/D137 | 0.28 | Q146/D150 | 0.23 | R48/D128 | 0.16 |
| I89/V126 | 0.45 | S8/N9 | 0.26 | N124/A85 | 0.21 | N124/A85 | 0.12 |
| A85/I80 | 0.38 | Y79/Q37 | 0.22 | P119/A116 | 0.21 | Q146/D150 | 0.12 |
| S35/D32 | 0.37 | H83/D41 | 0.22 | S117/T114 | 0.19 | S139/D137 | 0.12 |
| K139/D137 | 0.26 | P119/A116 | 0.17 | S139/D137 | 0.18 | S8/I46 | 0.11 |
| R48/Q90 | 0.21 | Q146/D150 | 0.16 | R48/D128 | 0.13 | Q37/Y79 | 0.11 |
| Q42/P39 | 0.20 | D90/V49 | 0.15 | D90/V49 | 0.12 | Y69/D66 | 0.08 |
| Top 30: | 18.79 |  | 17.20 |  | 15.56 |  | 14.61 |
| All Interactions: | 20.15 |  | 18.77 |  | 17.12 |  | 15.82 |

The hydrogen bond from Y77 to V53 is made possible by other aromatic residues in the vicinity subtly perturbing the relative orientation of the α_2_ and α_3_ helices. In *Po*Cel5, residue 123 is a tyrosine, which interacts with T78 to satisfy its hydrogen bonding requirements in an otherwise hydrophobic region of the protein (Fig. 4C). The effective Y123F mutation in the hybrids and *Te*Egl5A, coupled to N122 hydrogen bonding to S78, creates a hydrophobic pocket full of favorable π-stacking interactions (Fig. 4D). The stacking interactions maintain a favorable environment for Y79 without causing a helix rotation that would destroy the structure, in turn causing the carbonyl of V53 to have an alternate hydrogen bonding partner. These interactions together mean that the hydrophobic packing improves at this interface relative to the mesophile, a phenomena that has been noted in interaction clusters in other cellulases (36).

An alternative method of quantifying these hydrophobic interactions is to determine the number of contacts between different structural elements within the enzyme. The overall contact structure highlights the similar fold between all of the enzymes considered here (Fig. S4). However, the most revealing aspect of contact analysis occurs when comparing the number of contacts across different temperatures (Fig. 5). As expected, the number of observed contacts tends to decrease at high temperature due to the increased fluctuation computed previously (Fig. 3). The singular clear exception are the contacts between α_2_ and α_3_, which actually increase with temperature in the hybrids and *Te*Egl5A. The hydrophobic effect strengthens at higher temperature (37), driving the observed aromatic packing within the simulations.

**FIGURE 5.**
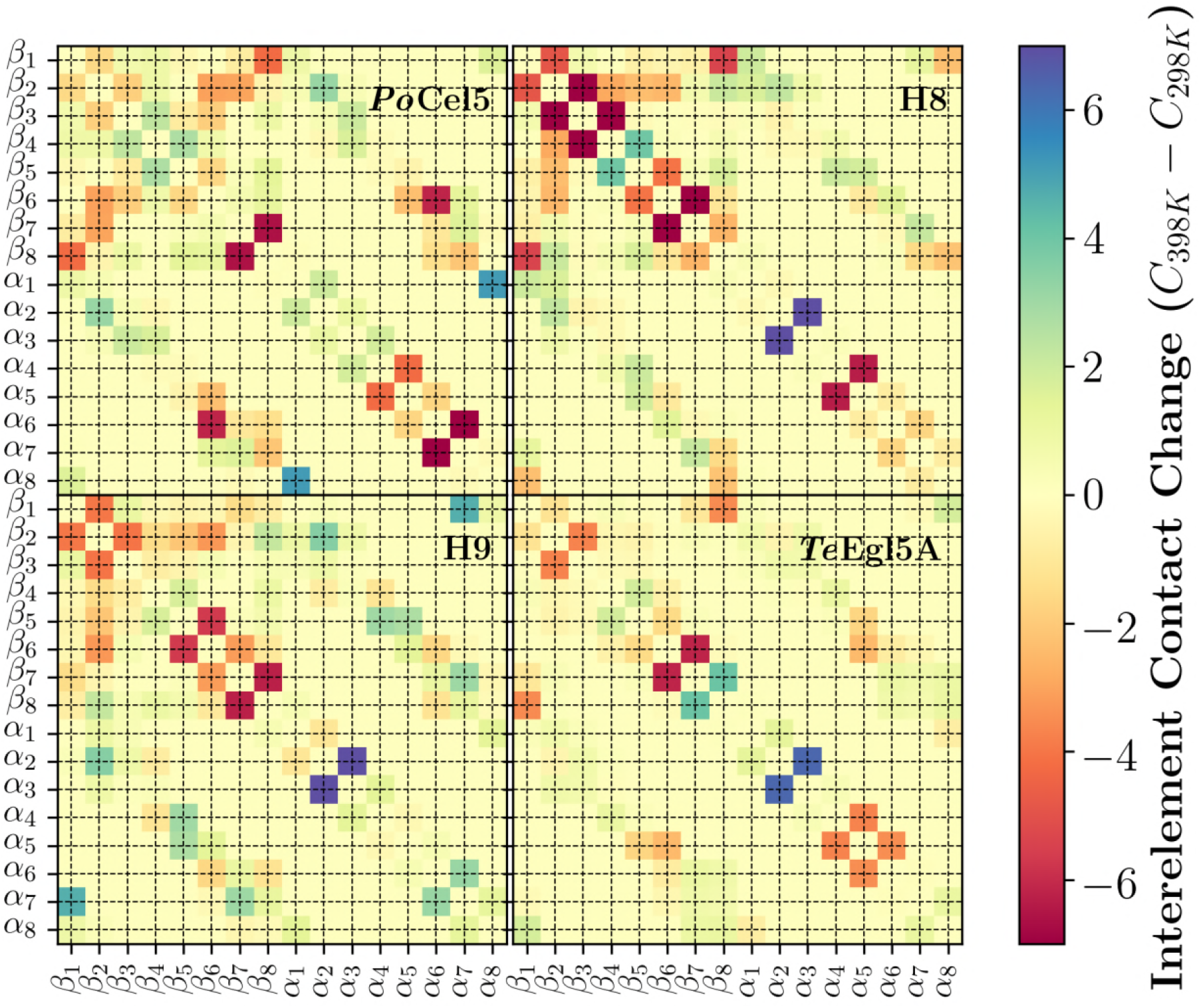
Atomic contact change between different structural elements in going from 298K to 398K. Each of the four enzymes studied is labeled in the upper right of each subpanel, with the colorscale defined on the far right. Contacts were defined between individual atom pairs that were within 5 Angstroms during simulation, and weighted according to distance. The weighting function used was 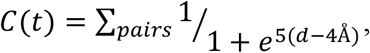, where d is the distance between individual atoms in the pair.

## DISCUSSION

Protein structure and function evolve through sequence changes, substitutions, duplications, insertions, and deletions, including rearrangement or recombination of short fragments as we did here, building on earlier work showing that folded hybrid domains can be generated by shuffling polypeptide segments (38). Based on prior experimental and structural studies suggesting that the common (βα)8 barrel or TIM barrel evolved from an ancestral half-barrel through a series of duplication, fusion, and diversification events (20, 39), including specific research in glycoside hydrolases (40, 41), our work here is in effect accelerated evolution with a particular design goal in mind. The research here is designed to combine the high substrate affinity of a mesophilic enzyme with the thermostability of a thermophilic enzyme to produce more efficient enzymes for specific substrates.

The key to the success of the improved enzyme variants was replacing the N-terminal half barrel, as the C-terminal half-barrel hybrid was not functional. This has precedence elsewhere in the literature for TIM barrel enzymes. In previous study, Prerna Sharma and coworkers reported that a stable and active chimera CelBCelCCA, which showed maximum activity at 70°C and was created by fusion the N-half barrel of the thermophilic CelB (maximum activity at 95°C) and C-half barrel of mesophilic CelCCA (maximum activity at 50°C) (40). Furthermore, Numata et al. constructed chimeric isopropylmalate dehydrogenases by connecting fragments from a thermophilic and a mesophilic parental enzyme. They found that the thermal stability of the chimeric enzymes was nearly proportional to the fraction of the sequence coming from the thermophilic enzyme, suggesting that amino acid residues contributing the thermal stability distribute themselves in the N-terminal half (42). Together with our results, this suggests that the N-terminal half barrel determines the thermostability of TIM barrel proteins, possibly by being the first part to unfold completely at the melting transition, although this was not observed over the short simulation timescales.

The observed high activity of two hybrid enzymes on specific substrates was also not guaranteed. Recombination of the segments between related enzymes often results in hybrids with diminished activity. For example, Hosseini-Mazinani and coworkers created 18 hybrid genes by substituting the coding region of the *P. vulgaris* β-lactamase gene with the equivalent portions from the RTEM-1 gene (43). Most of these hybrids produced inactive proteins, and a few hybrid enzymes had partial or trace activity. Even though the previously mentioned chimera CelBCelCCA displays hyperthermophile-like structural stability, the chimera activity is significant lower than those of the parental enzymes CelB or CelCCA (40). Similarly, in our research, half of the ten hybrids showed no activity even though they were constructed with similar methodology as the other five that were active, implying that more research will need to be done to determine what are the determinants of chimeric protein function.

To our surprise, two hybrid enzymes (H8 and H9) demonstrated increased the enzyme specific activity and catalytic efficiency using carboxymethylcellulose sodium (CMC-Na) as a substrate. Normally, mutants with increased stability often lose catalytic efficiency because flexibility is required for enzyme activity, whereas structural rigidity improves thermostability (32, 44). In contrast to this, both H8 and H9 showed improvement on thermostability and catalytic efficiency, which reduces the enzyme loading (and thereby the cost) required for production. We suspect that the systematic nature of the SCHEMA replacement and high structural homology of the TIM barrel proteins resulted in correctly positioned amino acids that were in the appropriate position for substrate binding and catalytic bond cleavage after functional domain recombination.

Parental enzymes that are sufficiently related lend themselves to the construction of hybrid proteins. Recombination segments or subdomain of proteins lies in the high similarity, thus the interaction required for the proper structure and function of hybrids will be retained. Then, recombinants are analyzed in an attempt to identify determinants responsible for parameters such as thermostability or activity (45). Based on the identity of the modules replaced in hybrids exhibiting enhanced thermostability, H8 and H9, and to some extent H6, the reasonable conclusion is that the interfaces between the β2, α2, β3, and α3 elements within *Po*Cel5 are stabilized through hybridization with *Te*Egl5A. In addition, with the results showing the mechanism behind the melting point improvement of the H8 and H9 hybrids relative to the progenitor *Po*Cel5, we can begin to speculate on the thermal denaturation mechanism of *Po*Cel5. Since the hybrid enzymes strengthened hydrophobic interactions between the second and third module to improve the melting point, this suggests that native *Po*Cel5 unfolding is initiated by water disrupting the packing between α_2_ and α_3_. One potential mechanism seen in simulation is T78 rotating away from its hydrogen bonding interaction with Y123 when it has the thermal energy to do so, drawing water to interact closely with Y123. In the H6, H8, and H9 hybrids, increasing the temperature strengthens the aromatic hydrophobic interactions at this same interface, raising the melting point until another structural element denatures and disrupts the structure.

The order of the relative melting points between H6, H8, and H9 can be used to inform the order of denaturation. Hybrid H6 has a lower melting point than H8 or H9 while sharing the thermophilic version of modules 2 and 3, but not module 1. This suggests that a structural element in module 1 unfolds first, likely α_1_ due to its greater accessibility to solution. After α_1_ unfolds, α_2_ would then not be packed against it, and may then drift away from α_3_, perturbing the packing between α_2_ and α_3_ and leading to enzyme unfolding. Our analysis finds no specific interactions with the remainder of the enzyme that would stabilize α_1_. Instead, our hypothesis is much simpler, and is motivated by sequence changes at the N-terminal end of α_1_. Residue 35 of *Te*Egl5A is a proline, compared with a tyrosine in *Po*Cel5. Prolines at the N-terminal end are known to raise the melting point of α-helices (46-48). Similarly residue 36 of *Te*Egl5A introduces an additional N-glycosylation site relative to *Po*Cel5. Glycosylation is also known to increase the thermostability of α-helices (49, 50). Together, these changes maintain the structure of α1 up to higher temperature, which in turn protects the packing of the α2 and α3 helices.

This does not, however, mean that α_1_ is the first helix to unfold. Even considering the short simulated timescales, we observed a loss of structure in α5 even at modest temperature, as evidenced by the high RMSF in that part of the protein (Fig. 3). It may be that this helix is natively in equilibrium between its folded and unfolded states, and that the adjacent residues were uniquely chosen to form compensatory interactions with the unfolded helix, preserving the overall structure. Such compensatory interactions would provide a mechanism for the melting point differential between H8 and H9, since only in H9 is the native α_4_-α_5_ interface is disrupted by the sequence changes made. However, another possible explanation is that the unfolding of α5 is a modelling artefact of adding in two additional alanines relative to the homologous crystallographic models.

In summary, through combinatorial swapping of sequence elements between a mesophilic and thermophilic cellulase, we created two enzymes with higher efficiency than either of the parental enzymes. These successful hybrid enzymes demonstrate increased thermostability relative to the mesophilic parent, while still less than that of a true thermophile. The high efficiency and improved thermostability may be useful for reducing enzyme loadings within industrial processes. Through the companion MD simulations performed at a ladder of temperatures, the interactions that lead to enhanced thermostability have been identified on the α2-α3 interface, which improves residue packing. Recapitulating the R52 interactions identified with surrounding anionic residues in the thermophilic *Te*Egl5A may be a further method of stabilizing the newly created hybrid enzymes.

## MATERIALS AND METHODS

### Strains, plasmids, and chemicals

The donor *P. opalus* CBS 125034 strain was cultivated at 28°C in an inducing medium containing 5 g/L (NH_4_)_2_SO_4_, 1 g/L KH_2_PO_4_, 0.5 g/L MgSO_4_·7H_2_O, 0.2 g/L CaCl_2_, 10 mg/L FeSO_4_·7H_2_O, 30 g/L corncob, 30 g/L soybean meal, and 30 g/L wheat bran. *Escherichia coli* Trans I-T1 (TransGen, Beijing, China) was used for gene cloning and construction of the hybrid enzymes. The vector pPIC9 and *Pichia pastoris* GS115 from Invitrogen (Carlsbad, CA) was used for enzyme expression. Yeast extract peptone dextrose (YPD), minimal dextrose (MD), buffered glycerol complex (BMGY), and buffered methanol complex (BMMY) were prepared according to the *Pichia* Expression Kit manual (Invitrogen). The FastPfu DNA polymerase from TransGen, restriction endonucleases from Fermentas (Burlington, Ontario, Canada), T4 DNA ligase from New England Biolabs (Hichin, UK), and BglII from TaKaRa (Kyoto, Japan) were purchased. The substrates carboxymethyl cellulose-sodium (CMC-Na), barley β-glucan, Avicel, laminarin, birchwood xylan, and locust bean gum were supplied by Sigma-Aldrich (St. Louis, MO). Lichenan was purchased from Megazyme (Wicklow, Ireland).

### Cloning and sequence analysis of the gene *Pocel5*

The genomic DNA of *P. opalus* CBS 125034 was extracted using a DNA isolation kit (Tiangen, Beijing, China). The total RNA was extracted, reverse transcribed, and used as a template for cDNA amplification as described by Zhao et al. (51). Using the genomic DNA of *P. opalus* CBS 125034 as a template, the core region of the cellulase-encoding gene *PoCel5* was amplified with a degenerate primer set GH5-F/GH5-R (Table S1), and its 5′- and 3′ - flanking regions were obtained by thermal asymmetric interlaced (TAIL)-PCR with four arbitrary degenerate primers from TaKaRa Genome Walking Kit and four nested specific primers (us-1, us-2, ds-1, and ds-2) (Table S1) designed based on the core region sequence of *Po*Cel5 (52). The PCR products were ligated with pEASY-T3 vector for sequencing and assembled to give the full-length *Po*Cel5 (GenBank accession number ARO48344). The expression primers *Pocel5-F/Pocel5-R* with EcoRI and NotI restriction sites (Table S1) were used to amplify the cDNA fragment coding for mature *Po*Cel5 without the putative signal peptide sequence. The PCR product was purified using the Gel Extraction Kit (Omega Bio-tek, Norcross, GA) and then ligated into the pPIC9 expression vector to produce the recombinant expression vector pPIC9-*Pocel5*.

The DNA and amino acid sequences were analyzed using the BLASTx and BLASTp programs (http://www.ncbi.nlm.nih.gov/BLAST/) (53), respectively. The introns, exons, and transcription initiation sites were predicted using the GENSCAN Web Server (http://genes.mit.edu/GENSCAN.html) (54). SignalP 3.0 was used to predict the signal peptide sequence (http://www.cbs.dtu.dk/services/SignalP/) (55). Sequence assembly and estimation of the molecular mass and *p*I of the mature peptide were performed using the Vector NTI Suite 10.0 software (Invitrogen).

### Production and purification of the recombinant *Po*Cel5

The recombinant plasmid pPIC9-*Pocel5* was linearized by BglII and transformed into *P. pastoris* GS115 competent cells by electroporation. Transformants were selected on plates of minimal dextrose medium at 30°C for 2 days. The positive clones were grown in shaker tubes containing 3 mL of BMGY at 30°C for 48 h, followed by cell collection and enzyme induction in 1.5 mL BMMY containing 0.5% methanol at 30°C for 72 h. The culture supernatant of each transformant was collected by centrifugation at 12,000 × *g* for 10 min at 4°C and examined by both SDS-PAGE and a cellulase activity assay. The positive transformants were also verified by colony PCR and sequencing. For scale-up cultivation, the transformant with highest cellulase activity was grown in 1-L Erlenmeyer flasks containing 400 mL of BMGY at 30°C for 48 h with an agitation rate of 200 rpm. Cells were harvested and resuspended in 200 mL of BMMY containing 0.5% (v/v) methanol for 48-h induction at 30°C.

The culture supernatants were collected by centrifugation at 12,000 × *g* for 10 min at 4°C, followed by ultrafiltration using a vivaflow 50 ultrafiltration membrane with a molecular weight cut-off of 10 kDa (Vivascience, Hannover, Germany). The crude enzyme was applied to HiTrap Q HP anion exchange column (Amersham Biosciences, Uppsala, Sweden) equilibrated with a 10 mM phosphate buffer of pH 7.5. A linear NaCl gradient of 0 to 1 M was used to elute the proteins. The apparent molecular mass and purity of purified recombinant *Po*Cel5 were estimated by SDS-PAGE. Endo-β-*N*-acetylglucosaminidase H (Endo H) from New England Biolabs was used to remove *N*-glycosylation according to the manufacturer’s instructions.

### Enzyme characterization

The cellulase activity was determined by using the dinitrosalicylic acid (DNS) method (56). Reactions containing 100 μL of properly diluted protein solution and 900 μL 1% (w/v) CMC-Na were incubated at pH 5.0 (100 mM citric acid-Na_2_HPO_4_) and 60°C for 10 min, followed by the addition of 1.5 mL DNS solution and 5 min in a boiling water bath. When the reactions cooled to room temperature, the absorbance at 540 nm was measured. The standard curve for calibrating the enzyme activity was determined by 0.25–3.5 μmol glucose. One unit of enzyme activity was defined as the amount of enzyme required to release 1 μmol of reducing sugars per min. Specific activity was defined as the enzymatic units per milligram protein.

The optimal pH for *Po*Cel5 was determined at 60°C in 100 mM citric acid-Na_2_HPO_4_ buffer ranges of 3.0–8.0. The optimal temperature was determined at pH 5.0 in the temperature range of 50–90°C. The pH stability was determined by measuring the residual activity at pH 5.0 (100 mM citric acid-Na_2_HPO_4_) and 60°C for 10 min after 1-h incubation at pH 3.0–12.0 and 37°C without CMC-Na. Thermostability was investigated after incubation of the samples at pH 5.0 and 70°C for different periods of time. Residual activity was measured as described above.

Substrate specificity of *Po*Cel5 was determined by using 1% CMC-Na, barley β-glucan, lichenan, birchwood xylan, Avicel, laminarin, or 0.5% locust bean gum as the substrate. The kinetic parameters *K*_m_, *k*_cat_, *V*_max_, and *k*_cat_/*K*_m_ were estimated from the *Po*Cel5 activities at pH 5.0 and 60°C for 5 min by using 1–10 mg/mL CMC-Na as the substrate. GraphPad Prism 6.0 (http://www.graphpad.com/scientific-software/prism/) was used to calculate the values by using the Lineweaver-Burk plot.

### Design and construction of the hybrid enzymes

*Po*Cel5 shared 51% and 47% sequence identity with the thermophilic endoglucanase *Te*Egl5A (GenBank accession number KF680302) and its N-terminal (βα)_4_ modules (Fig.1), respectively. By using the fusion protein method, the N-terminal (βα)_1–4_ module(s) or the C-terminal (βα)_5–8_ modules of *Te*Egl5A were introduced into the corresponding parts of Pocel5 (Table 1), and a total of 10 hybrid enzymes were constructed. To further verify the functional roles of the N-terminal sequence of *Te*Egl5A in another GH5 cellulase, *An*Cel5 (GenBank accession number AAG50051) from *Aspergillus niger* (11) sharing 56% identity with *Po*Cel5 and 67% identity with *Te*Egl5A was selected, and hybrid enzymes were constructed by replacing the N-terminal (βα)_1–3_ and (βα)_1–4_ modules, analogous to hybrids H8 and H9 with *Po*Cel5.

All the fusion proteins were obtained by a two-step overlap extension PCR (Fig. S1). In the first-step PCR, parallel reactions were performed to amplify the objective DNA fragments using the recombinant plasmids pPIC9-*Teegl5A*, pPIC9-*Pocel5* and pPIC9-*Ancel5* as templates and primers. The specific recombinations are listed in Table S1. The second-step PCR was used to amplify the final DNA products with the first-step PCR products as templates and primer sets F/A and R/B/D (Table S1). The 50-μL PCR mixture contained 1 μL of each primer, 1 μL of Fastpfu Fly DNA Polymerase (TransStart, Beijing, China), 5 μL of dNTPs, 10 μL of Fastpfu Fly buffer, 3 μL of MgSO_4_, 1 μL of template DNA, and 28 μL of ddH_2_O. The PCR protocol contained an initial denaturation at 95°C for 5 min, followed by 30 cycles of 94°C for 30 s, annealing at 60°C for 30 s, and elongation at 72°C for 60 s, with a final extension at 72°C for 10 min. The PCR products were purified, digested with EcoRI and NotI, ligated into the pPIC9 expression vector, and sequenced. The production, purification, and characterization of hybrid enzymes followed the same procedures of recombinant *Po*Cel5A.

### Thermal stability analysis

For short-term thermostability analysis, the purified recombinant wild-type and hybrid enzymes were incubated at 70°C and/or 80°C and optimal pH for 1-h without substrate. Samples were taken at specific time points for the cellulase activity assay under optimal conditions of each enzyme. Three thermodynamic parameters, *T_50_*, *t*_1/2_, and *T_m_*, were used to compare the thermal properties of *Po*Cel5, *An*Cel5, and their hybrid enzymes. *T_50_* was defined as the temperature at which a 30-min incubation caused the protein (0.1 mg/mL) to lose 50% activity, while *t*_1/2_ was defined as the half-life of an enzyme at 55°C (for *Po*Cel5 and its hybrid enzymes) and 70°C (for *An*Cel5 and its hybrid enzymes). Differential scanning calorimetry (DSC) was used to determine the *T_m_* values. The Nano DSC (TA Instruments) was run at a heating and scanning rate of 1°C/min over a temperature range of 30 to 90°C. Each sample contained 0.25 mg/mL protein in 10 mM citric acid-Na_2_HPO_4_ buffer (pH 7.5). The test was repeated at least twice.

### Molecular dynamics (MD) simulation

To determine the specific molecular interactions underlying the observed improvement in thermostability, a series of equilibrium classical molecular dynamics simulations of modelled GH5 structures were performed using NAMD 2.12 (57). Four GH5 models were constructed, one for each of the *Po*Cel5 and *Te*Egl5A parent enzymes as well as for the H8 and H9 chimeras. The creation of the four homology models used MODELLER 9.19 (58), using PDB structures 5I78 from *Aspergillus niger* (11), 1H1N from *Thermoascus aurantiacus* (59), and 5L9C from *Penicillium verruculosum* as the structural templates for system construction. These templates all have greater than 50% sequence identity with each of the four GH5 sequences considered (Fig. 1), which is typically indicative of strong structural homology (58). Protonation states consistent with the optimal activity pH (5.0) for each GH5 model were determined using PROPKA 3.1 (33, 60). Since some of the PDB reference structures feature N-glycosylations (11), which are commonly found in other glycoside hydrolases (2) in addition to our direct experimental evidence for glycosylations on our enzymes, between 1 and 3 basic N-glycosylations were added where appropriate to accessible asparagine residues on each model using the GlyProt webserver (61). Specifically, *Po*Cel5 has glycosylations on Asn23, Asn64, and Asn76. *Te*Egl5A has four glycosylations, on Asn36, Asn 190, Asn219, and Asn267. H8 and H9 each have a single glycosylation at Asn36. Other asparagine residues were determined to be inaccessible to solution. Each of the four complete GH5 models was solvated in a water cube with 80 Å sides using the SOLVATE plugin of VMD (62).

Following construction, each of the four GH5 models was simulated for 10ns where the alpha carbons were harmonically restrained to their initial positions using a 1 kcal mol^-1^ Å^-2^ force constant. This equilibration step permits the placed water molecules and the modelled side chains to find their preferred orientations and rotameric states given the backbone structure. The equilibration was performed in a constant pressure and temperature ensemble, using a Langevin piston (63) as the barostat to maintain 1 atm of isotropic pressure and a Langevin thermostat (64) set to 298K with a 1ps^-1^ coupling coefficient. The CHARMM36 (65, 66) force field was used for protein components, together with the CHARMM36 carbohydrate force field for the glycosylations (67, 68) and the TIP3 water model (69). Non-bonded terms of the energy function were cut off at 12 Å after a 10 Å switching distance. Long range electrostatic forces were determined using the particle mesh Ewald method (70, 71) with a 1.2 Å grid spacing.

The end state after this short equilibration was then used as the starting point for all subsequent simulations. For each model, simulations were carried out at three different temperatures, 298K (25°C), 343K (70°C, near the melting point), and 398K (125°C). To minimize the impact of the stochastic nature of these simulations on the final conclusions, each combination of GH5 model and temperature was simulated five times for 200ns each. The aggregate 4 microseconds of trajectory were then analyzed using a python-enabled build of VMD (62), using its built-in hydrogen bond analysis tools. Atomic contacts between structural elements were computed using a weighted contacts formula (72, 73), which were used to identify hydrophobic interactions that are otherwise difficult to quantify.

## Acknowledgements

This work was supported by the National Natural Science Foundation of China (no. 31572446), the Fundamental Research Funds for Central Non-profit Scientific Institution (no. Y2017JC31), and the National Chicken Industry Technology System of China (no. CARS-41). GTB thanks the U.S. Department of Energy (DOE) Energy Efficiency and Renewable Energy (EERE) BioEnergy Technologies Office (BETO) for funding under Contract No. DE-AC36-08GO28308 the National Renewable Energy Laboratory (NREL). JVV was supported by the NREL Director’s Fellowship funded by the Laboratory Directed Research and Development (LDRD) program. This work used computing resources provided by the Extreme Science and Engineering Discovery Environment (XSEDE) (74), which is supported by National Science Foundation grant number ACI-1548562. Specifically, the allocation to GTB (TG-MCB090159) was used on Stampede2, which is hosted by the Texas Advanced Computing Center (TACC) at the University of Texas at Austin. The U.S. Government retains and the publisher, by accepting the article for publication, acknowledges that the U.S. Government retains a nonexclusive, paid up, irrevocable, worldwide license to publish or reproduce the published form of this work, or allow others to do so, for U.S. Government purposes.

## Author contributions

FZ performed the experiments and wrote the manuscript. JVV set up and analyzed the molecular dynamics trajectories and wrote the manuscript. JZ, YW, TT, XW and XM helped analyze the data and revised the manuscript. BY, GTB and HL revised the manuscript. All authors read and approved the final manuscript.

